# The dynamics of decision making and action during active sampling

**DOI:** 10.1101/2021.03.10.434801

**Authors:** Duygu Ozbagci, Ruben Moreno-Bote, Salvador Soto-Faraco

**Affiliations:** Center for Brain and Cognition and Department of Information and Communications Technologies, Pompeu Fabra University, Barcelona, Spain; Institut Català de Recerca I Estudis Avançats (ICREA), Barcelona, Spain

**Author notes:** Corresponding Autor.

**Keywords:** decision-making, embodied cognition, mouse tracking, action dynamics, motor action

## Abstract

Embodied Cognition Theories (ECTs) propose that the decision process continues to unfold during the execution of choice actions, and its outcome manifests itself in these actions. Scenarios where actions not only express choice but also help sample information can provide a valuable test of this framework. Remarkably almost no studies so far have addressed this scenario. Here, we present a study testing just this paradigmatic situation with humans. On each trial, subjects categorized a central object image, blurred to different extents (2AFC task) by moving a cursor toward the left or right of the display. Upward cursor movements, orthogonal with respect to choice options, reduced the image blur and could be freely used to actively sample information. Thus, actions for decision and actions for sampling were made orthogonal to each other. We analyzed response trajectories to test a central prediction of ECTs; whether information-sampling movements co-occurred with the ongoing decision process. Trajectory data revealed were bimodally distributed, with one kind being direct towards one response option (non-sampling trials), and the other kind containing an initial upward component before veering off towards an option (sampling trials). This implies that there was an initial decision at the early stage of a trial whether to sample information or not. Importantly, the trajectories in sampling trials were not purely upward, but rather had a significant horizontal deviation that was visible early on in the movement. This result suggests that movements to sample information exhibit an online interaction with the decision process. The finding that decision processes interact with actions to sample information supports the ECT under novel, ecologically relevant constrains.

## 1. Introduction

The classical view of decision-making was founded on the idea that action is executed after a decision has been made, in a serial fashion (e.g., Newell & Simon, 1972). This idea assumes a clear temporal and functional separation between decision making and the motor processes that implement that decision. However, later behavioural studies challenged this serial view and proposed that decision does not have to finish before the movement execution process begins, *de facto* introducing the parallel view of decision making (e.g., Ghez, et. al., 1997 & McKinstry, et. al., 2008). This parallel view states that there is an ongoing information flow from decision to action systems well before the decision process has been fully completed. According to this view, not only action may start before a decision is reached, but movements may be updated online based on newly acquired evidence (Coles, et. al., 1985).

To investigate the putative interaction between action and decision as it unfolds in time, some studies have used tasks which require continuous control of action. To this end, decision-making tasks incorporated movement tracking for responses executed on devices like a joystick, a robotic handle, a computer mouse, or freely with a hand reaching movement (Resulaj, et. al., 2009, Burk, et.al., 2014, Barca & Pezzulo, 2012; Song & Nakayama, 2008). Since these responses have a wide temporal and spatial span, they make it possible to study, and compare the dynamics of the movement during the decision-making process.

A typical finding that emerges from continuous movement paradigms when subjects must move toward one out of two alternative targets, is the prevalence of movement trajectories that are not perfectly direct to the chosen target (Song & Nakayama, 2008). These findings have shown that the initial phase of the response movement weighs in the paths to the two possible targets, maintaining a compromise which is later resolved by diversion of the trajectory committing to one of the targets (Chapman, et. al., 2010 & Gallivan, et. al., 2011). These averaged movement trajectories are interpreted as a case of movement being planned and executed online during the deliberation process and more importantly, that there is a continuous crosstalk between these two processes (Cisek & Pastor-Bernier, 2014, Marcos, et. al., 2015). An exacerbated expression of this are changes of mind, trials in which the subject’s response movement starts off toward one target but corrects on-the-fly toward the alternative target (Burk, et. al., 2014). In general, these findings motivated the parallel view of decision making, which focuses on the ongoing one-way flow of information from action to decision.

Although the parallel view of decision making assumes a richer interaction between action and decision than the sequential view, it only accounts for the forward influence from decision to action. However, there is evidence for backward influence from action on decision as well. For example, Burk, et. al. (2014) showed that when the spatial distance between two response options is large, subjects make less changes of mind than when the distance between targets is shorter. This means that action costs are considered and influence the outcome of the decision process. Similarly, Cos, et. al. (2011) found that the amount of effort required to perform the response action biased performance in a perceptual decision-making task.

We can frame the evidence mentioned above under Embodied Cognition Theories (ECT) of decision-making, whose common characteristic is the influence of action dynamics on decision. Indeed, drawing connections between motor processes and decision making has a conceptual grounding on the wider framework of sensorimotor and embodied views in cognitive sciences (Clark, 1999, O’Regan & Noe, 2001, Barsalou, 2008), a general conceptual shift that has pervaded recent views in decision-making. One clear example is Lepora & Pezzulo’s (2015) Embodied Choice Model. The model proposes a two-way online interaction between motor actions and decision processes and that this interaction allows for a fast update of movement and decision processes. A typical argument by example often used to support this view is that, in nature, animals must move about (their body and/or sensory epithelia) to be able to perceive information that is necessary for making choices and planning actions (see Lepora & Pezzulo, 2015). To use the information gained through movement though, there needs to be a backward flow of information from action-related motor processes to decision making.

Despite the logical emphasis that embodied views make on information sampling movements, this notion has not been implemented in experimental tasks to support the ECT. In fact, in most of these decision making tasks, the stimulus information is available all at once and static, without any dependency upon the participant’s movement (Lepora & Pezzulo, 2015, Barca & Pezzulo, 2012, Hudson, et. al., 2007, Marcos, et. al., 2013). The interactions in these types of tasks have been illustrated in Figure 1a. Because the actions performed to report a choice are inconsequential to the inflow of information used to reach that decision, these tasks cannot capture all possible interactions between action and decision proposed by ECTs. Therefore, there is a need for tasks that can reveal the two relevant aspects of actions to identify the potential interplay between motor and decision processes. This interplay, which has motivated the task used here to test decision making under ECT, is illustrated in Figure 1b. Here, we assume that there are two types of action plans which are critical in an embodied decision-making scenario, the ones necessary for response itself, and the ones necessary for information sampling. Both of them interact with the decision process, and mediate both feed-forward and feedback interactions.

**Figure 1.**
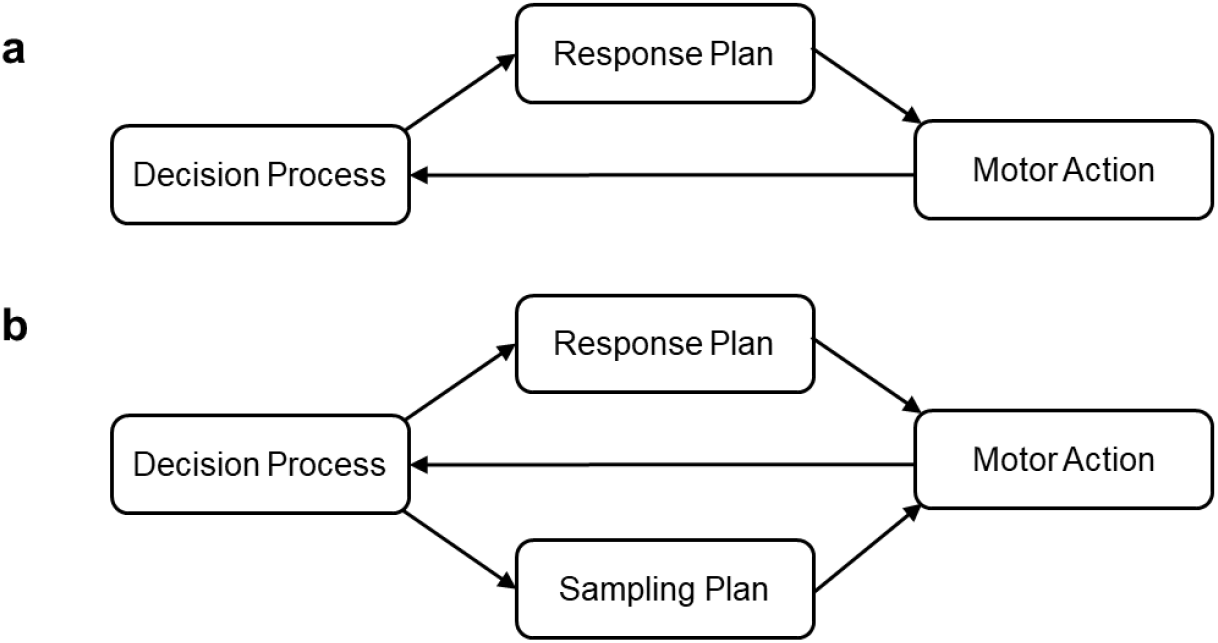
Interactions between motor action and decision in tasks without **(a)** and with **(b)** active information sampling. **a.** In classical tasks (see text) decision process feeds the response plan which gets executed with a motor action. While the action continues, the output of the action feeds back into the decision process. This is not a fully embodied scenario, since actions do not bring an information change. **b.** In a fully embodied scenario considered here, two different action plans, for sampling and for responding, are allowed to unfold in parallel. The decision process has a feedforward influence on motor output, whereas sampling influences decision via feedback from the motor action. In contrast to panel **(a)**, the executed motor action implements both responding and sampling of information.

In conclusion, we believe that the generality of the interplay between decision and action, and by proxy, of the embodied decision framework, have not yet been tested in all its critical components. In the present study, we aim at testing the ECT’s predictions with a task in which information accrual depends on the subject’s actions. We have developed a novel mouse-tracking task in which action is necessary both to sample information and to indicate the decision. To be able to single out one from the other, movements directed to sample information and movements to respond have been made orthogonal. That is, it is possible for the subjects to accumulate all the information first and then make the choice, make a choice at once without any accumulation of information, or anything in between. Although sampling and response actions have orthogonal dynamics, one critical aspect of the task is that both action plans are executed via same effector, so that the final motor output must synthesise the two plans if they are to co-occur, as the theory predicts.

Our hypothesis, derived from the ECT (Lepora & Pezzulo, 2015, Cisek & Pastor-Bernier, 2014), is that the movements related to the decision-making process and the movements related to accumulation of evidence to reach that decision are subject to significant online interaction. We first show that, in our task, movements depend on the amount of available information such that participants move to sample information when needed. Second, we demonstrate that the decision-making process transpires even in the initial phases of the information sampling movements, so that trajectories are biased towards one (usually the chosen) target much before all the information has been gathered. These results do not only suggest that the decision-making process pervades information sampling actions, but also that decision, actions and information sampling are orchestrated in parallel, and not in a strictly sequential fashion.

## 2. Methods

### 2.1. Participants

Twenty-one voluntary participants joined the experiment (13 women, 9 men, average age 23.5 years). Participants were recruited from the database of the Center for Brain & Cognition (University Pompeu Fabra) and were paid 10 euros per hour in exchange for their participation. They were all right-handed and had normal or corrected to normal vision with no reported history of motor problems related to upper limbs. Before proceeding with the experiment, all subjects read and signed an informed consent form. The experimental protocol was approved by the ethics committee CEIC Parc de Salut Mar, Universitat Pompeu Fabra. Before conducting the hypothesis-driven data analyses, we excluded data from two subjects whose accuracy was below 75%.

### 2.2. Experimental setup

Participants were asked to perform a visual object categorization between “edible” vs “non-edible” in a two-alternative forced choice (2AFC) paradigm. We used 63 edible and 63 non-edible object images from the Amsterdam Library of Object Images (Geusobroek, et. al., 2005), and each of them was presented only once to each participant, obtaining a total of 126 different trials per participant. To control for possible effects of colour cues, we used grey-filtered versions of the images. Stimulus display and the task were programmed with MATLAB, PsychToolBox (Brainard, 1997). Visual stimuli were presented on a Cambridge Research Systems, Display++ monitor (1920 × 1080 pixels, 32’’, 100 Hz refresh rate). Responses were recoded through a computer mouse (HP USB Optical Scroll Mouse), and the cursor location was recorded at 100 Hz (at every display refresh frame). The participant’s task involved moving the cursor from a home position at the bottom centre of the display to the right or left response areas, depending on the choice regarding the image presented at the top centre (locations and other details are described below).

For each subject, the total of 126 trials were divided, randomly and equiprobably into three different movement-to-visibility conditions: No Blur (NB), Low Blur (LB) and High Blur (HB). In the NB condition, the images were fully visible (without any blur) from the beginning of the trial, and therefore visibility was not contingent on action. For the other two conditions, in order to implement movement-dependent updating of information, we manipulated the visibility of the object images as a function of mouse position. We used a dynamic filter mask over the image to blur the image. The filter convolved each pixel with the neighbouring pixels with a Gaussian kernel with standard deviation (sd) proportional to the vertical distance between current cursor position and the target image at the top centre of the display, denoted *d*_*v*_ (measured in pixels). In the LB condition, the Gaussian mask had sd = *d_v_*/120, whereas in the HB condition the Gaussian mask had sd = *d*_*ν*_/60. This effectively made blur (hence, image visibility) depend on the participants, movement, so that moving upward de-blurred the target image (i.e., the shorter the vertical distance to target, the smaller *d*_*ν*_, and hence the lower the sd and the higher the visibility). The difference between the two blur conditions was the gain in visibility as a function of distance.

### 2.3. Procedure

Each subject completed the task in a darkened, sound-attenuated laboratory room. Before each trial started, the subject moved the mouse cursor to the bottom-centre home area (height = 10 × width = 15 pixels, centre x, y coordinates: 960,1075 pixels). The trial began with the image (265 ×192 pixels) appearing at the top-centre of the monitor (x- coordinates: 827 to 1092, y-coordinates: 0 to 192 pixels). As soon as the image appeared, the subject was free to move the mouse to indicate her choice by reaching to, and clicking on, one of two response areas, left or right side of the display, within 2000 ms (Figure 2). The rectangular response areas, covering the leftmost and rightmost 23% of the display, were indicated by two vertical lines along the screen sides (x coordinates: 440 and 1480 pixels, respectively; see, Figure 2). For half of the participants, edible was attributed to the left response area and non-edible to the right. In the other half, it was reversed. Response deadline was 2000 ms, after which the subject missed the trial. Each trial took the whole 2000 ms, independently of the response time. After a trial ended, the participant needed to move the cursor back to the bottom-centre home location for the next trial to begin. The inter-trial interval was 2000 ms, which also served as a fixation screen. Trials from all three conditions (NB, LB, HB) were interleaved randomly throughout the experiment. Hence, for efficient responding, participants could not fall back on a pre-defined strategy based on visibility prior to the start of the trial.

**Figure 2.**
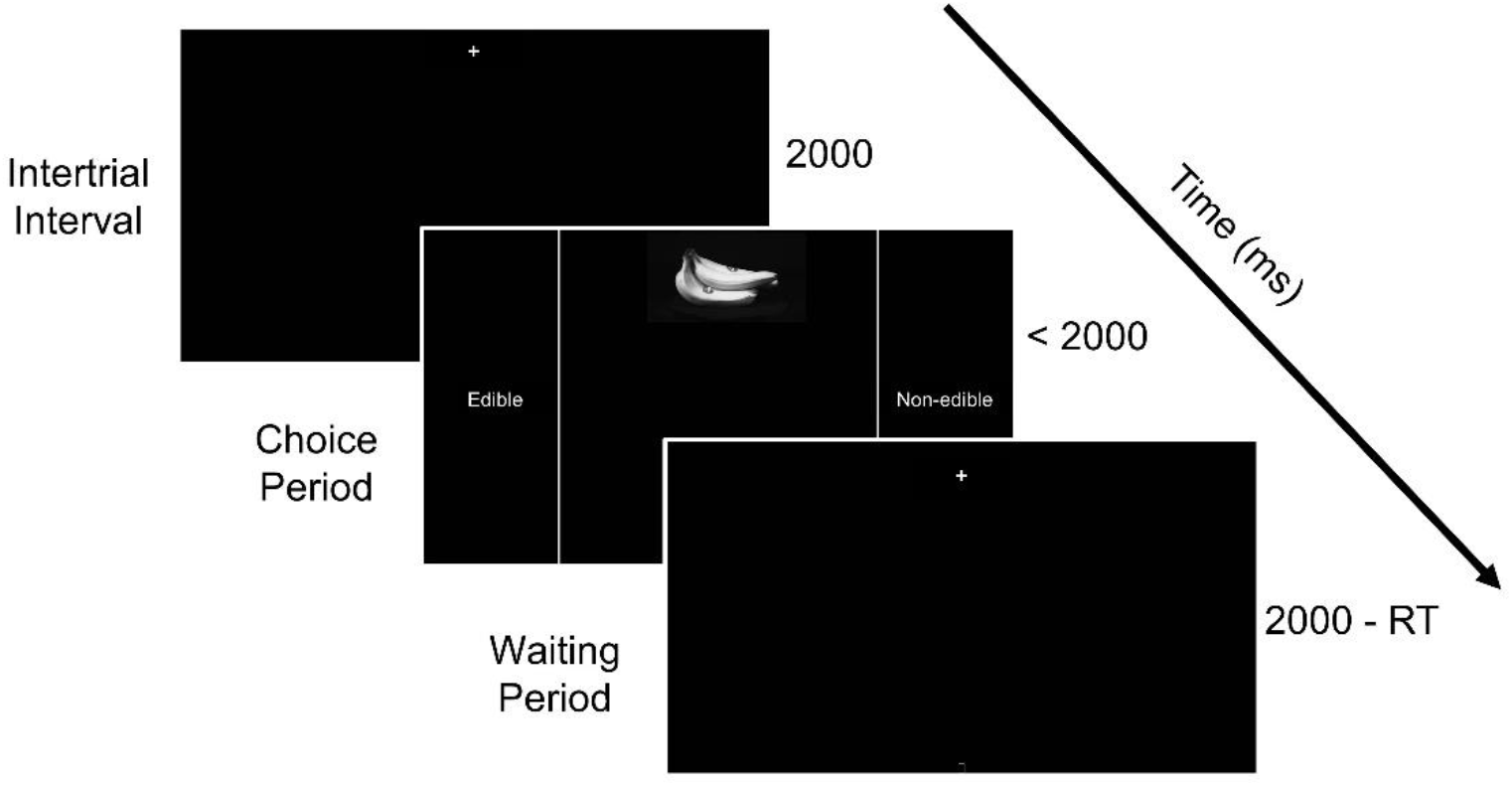
Schematic illustration of a trial sequence. Each trial was preceded by a 2000ms inter-trial interval displaying a fixation cross. Then, the stimulus and the choices were presented on the screen until response, with a deadline of 2000ms. Response areas, left and right of the display, are denoted by straight vertical lines. All trials were equated to the same duration, 2000ms by adding a waiting time if necessary. RT = reaction time.

Because the response areas covered both lateral sides of the display, the decision movement could vary in terms of the vertical extent of the trajectory, including direct horizontal movements from the home location to the response area. As said earlier, in the blurred image (LB and HB) conditions, the image blur decreased as the mouse moved upward. Therefore, when the image did not contain sufficient information, the participant needed to move in the vertical direction in order to gather evidence. Because of the response deadline (2000 ms), moving upward had a cost (i.e., took time off the available response time). Therefore, moving upward is never an optimal strategy if it is not necessary to sample evidence.

## 3. Results

In our task, characterizing information sampling and response components of the subjects’ action boils down to the analysis of heights and angles of the response trajectories (some example trajectories are shown in Figure 3). Firstly, we inspected the trajectory height, denoted *h*, which was calculated by measuring the vertical distance (in pixels) between the starting point and the highest point of the trajectory (Figure 3a). Second, we analysed the initial angle of trajectories, denoted α, which was defined as the angle described by an imaginary straight line connecting the starting point with the point at one-third of the length of the trajectory (cyan dashed line in Figure 3a), with respect to the vertical midline. It is important to note that, although correct targets were randomly assigned left or right sides during the task, for analyses we realigned the correct choice to positive angles.

**Figure 3.**
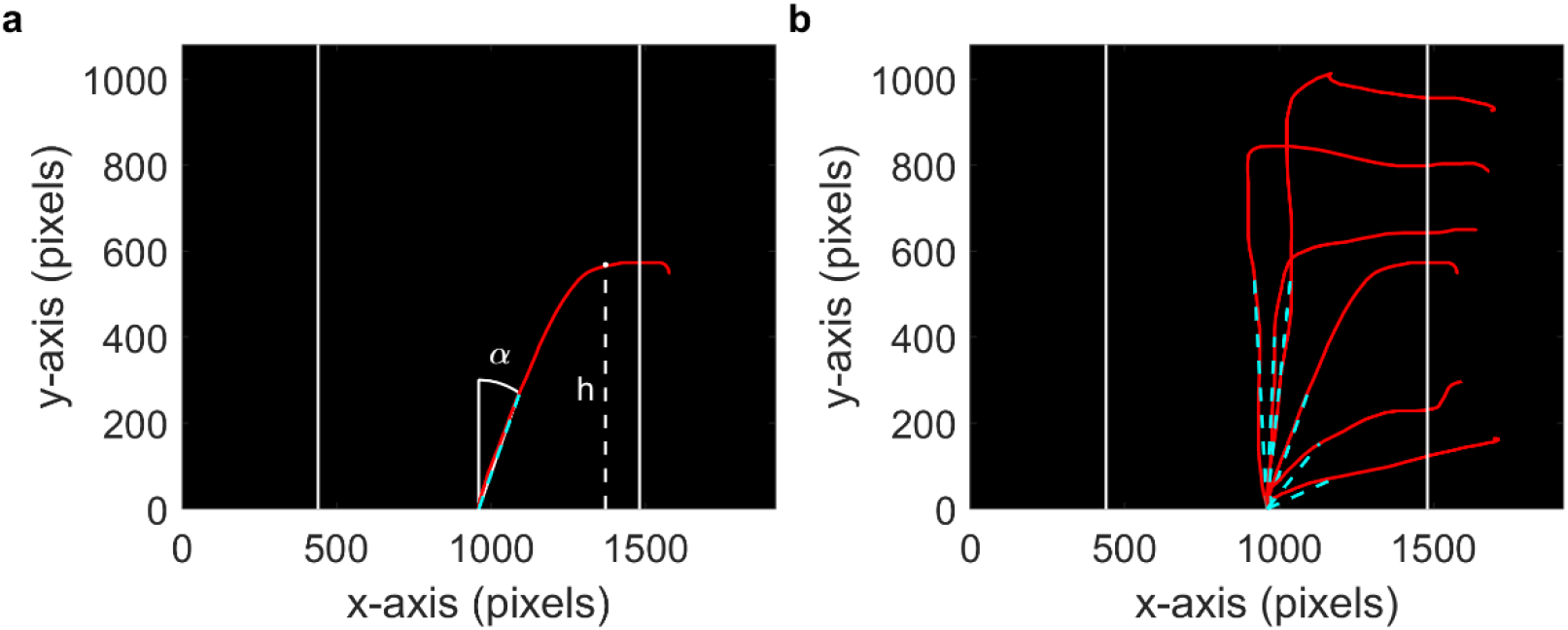
**a.** An example of one mouse trajectory (red line) on the experimental display. Response areas are indicated to the participants by the solid vertical lines on the left and right sides. The white dashed line indicates the height *h* of the trajectory. The cyan dashed line that joints the origin with the point of the trajectory that lies at one third of its total length serves to calculate the initial angle α of the trajectory with respect to vertical. Positive angles are defined to be in the direction of the correct target, whose location could occur randomly on either side. **b.** Examples of trajectories for several individual trials, with the same conventions described in a.

We preregistered this study and we first report the analysis that were planned prior to data collection (see, https://osf.io/3ysah/). We also performed follow-up analyses that have been planned after the pre-registration process, as these reveal important characteristics of the data. Throughout the results section we report statistical tests according to the frequentist approach (the analogous Bayesian analyses are reported in the Supplementary Table S1). Both analyses lead to the same conclusions. We included only the trajectories of correct trials into the analyses reported below (average 110 correct trials, >87%, out of 126 total per subject, range103-123).

### 3.1. Movement-dependent information sampling

If participants gather information as needed, their trajectories should reach higher when the image is blurred. We therefore tested whether trajectories in blur trials reached higher than trajectories in the no blur trials. As expected, trajectories in the two blur conditions were higher than in the no blur condition, since information sampling was unnecessary in the latter (right tail paired-samples t-tests, t(17) = 6.53, p <0.001, Cohen’s d = 1.54; t(17) = 7.03, p<0.001, Cohen’s d = 1.66, for the comparison of NB with LB and HB, respectively). However, even in the NB conditions trajectories had some vertical component (mean = 368.3 pixels, sd = 231.9), possibly due to biophysical motor constrains. To eliminate the height differences that are present in the trajectories but unrelated to information gathering, we subtracted the average height in NB condition from LB and HB trajectory heights in each individual’s data. Results (Figure 4a) showed that trajectories in HB trials were about 27% higher than in LB trials (mean = 315.7, sd = 190.4, vs 229.3, sd = 148.8, respectively; right tail paired-samples t-test, t(17) = 5.39, p<0.001, Cohen’s d =1.27).

**Figure 4.**
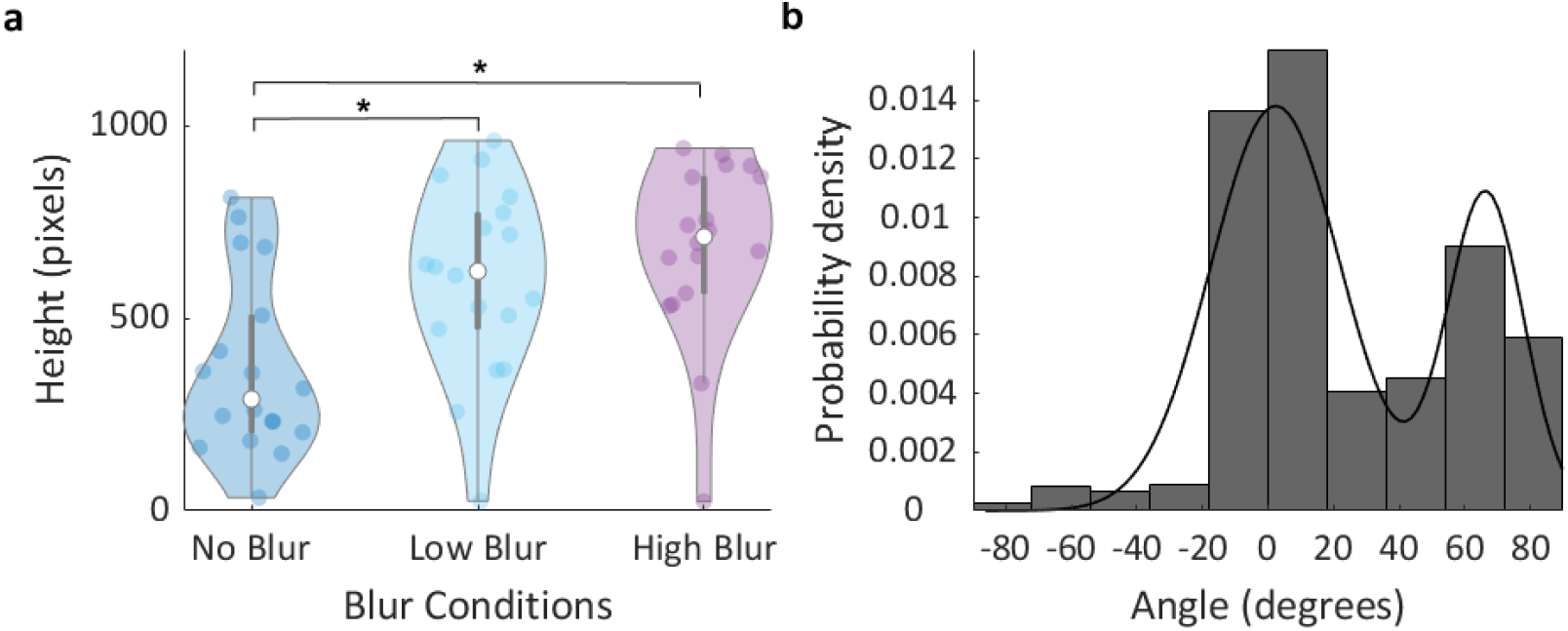
**a.** Height of trial trajectories for NB, LB and HB conditions. Each colored dot represents individual means for the corresponding condition. White dots represent the group median for the condition and the grey lines represent the inter-quartile range. **b.** Probability density of the initial angles of the trajectories across participants. The solid black line corresponds to the Gaussian mixture model (with 2 components) fit to the distribution (model with 2 components AIC = 19105 < model with 1 component AIC = 19753). Angle 0° corresponds to straight vertical upwards movement, i.e., with no horizontal component. Positive angles correspond to correct target direction.

### 3.2. Interplay between decision and action

#### Bimodality of trajectories

A central prediction of ECT is that movements should reflect the decision-making process throughout, such that the trajectories should show early on a bias towards the finally chosen target. We tested this prediction by studying the initial angles of the trajectories (Figure 4b). We found that their distribution is bimodal (Hartigan’s Dip Test, p-value<0.001; Gaussian mixture model better fit with 2 components, Akaike Information Criterion (AIC) = 19105 than the model with 1 component, AIC = 19753), with a central lobe peaking at angle 2.3°, a lateral lobe peaking at 66.4° (vertical midline corresponds to an initial angle of 0° and positive angles corresponding to directions to the correct target), and the separation between the two lobes being 43.52 °. This bimodality and the cut-off point allowed us to classify trajectories as sampling or non-sampling trajectories, depending on whether the initial angle is closer to the central or the lateral peak, respectively. We checked bimodality in the distribution of trajectory angles for each subject individually (see Figure S1) and found that 9 out of 18 subjects showed significant bimodality in the distribution in trajectory angles.

The presence of two types of trajectories is observed in each blur condition separately (Figure S2). While there is a large fraction of non-sampling trajectories in the NB condition (corresponding to the lateral lobe of the bimodal distribution; q = 0.62, X^2^(1, N = 635) = 39.86, p<0.001), surprisingly in the HB condition there were also non-sampling trajectories (q =0.14 binomial test p<.05). The presence of sampling and non-sampling trajectories across all blur conditions suggests that participants made an initial choice about whether or not to gather information, as supported by an analysis that shows that trajectories classified as non-sampling had a much smaller height than sampling trajectories (right tail two-sample t-test, t(1988) = 48.72, p<.001). Thus, non-sampling trajectories simply reflect a ballistic movement towards the chosen target that emanates from an initial decision, with little information gathering or ongoing decision-process throughout.

#### Angle analysis of sampling trajectories

Thus, given the initial decision and the ensuing existence of two different types of trajectories, a direct test of the prediction of ECT requires examining the sampling trajectories alone. These trajectories correspond to the central peak of the distribution in Fig. 4b. As the initial angles of these trials are close to zero (vertical), trajectories mostly depart vertically from the home position with the aim of gathering information to guide the final choice. However, in addition to the prominent vertical component, the initial steps of the trajectory were biased towards the chosen target, as the initial angle was significantly larger than zero in both LB and HB conditions (right tail one sample t-tests, t(16) =4.58, p<.001 and t(16) = 3.41, p = .002, respectively). This result strongly supports the notion that the decision-making process transpires into the movement even when participants felt urged to actively sample information.

One might argue that some trials in the analysis above might have been misclassified non-sampling trials, given the partial overlap of the two lobes of the bimodal distribution of angles. This could introduce some biases towards positive angles. To control for this possible confound we used a simpler analysis limited to LB and HB trials only (where participants are urged to sample information) that does not rely on trial classification. In this analysis we calculated average angle in incremental ranges of angles (symmetric around 0°) from ±1° to ±30°, in steps of one degree (Figure 5a). We found that the average angle was significantly larger than zero in all the ranges larger than ±14° (right tail t-tests, p <.05, see Figure 6a). Angles in the range ±14° and ±20° are well inside the central peak of the bimodal distribution, as described above, and therefore can be reliably classified as sampling trajectories (trajectories with such small initial angle very unlikely correspond to trials where the decision maker already made a choice about where to move). In sum, this new analysis shows trajectories whose initial angles lie within a small range of angles symmetric around zero already show a significant positive bias towards the chosen target. This result further supports the notion that the ongoing decision-making process transpires into the movement even when not all information necessary to solve the task has been gathered.

**Figure. 5.**
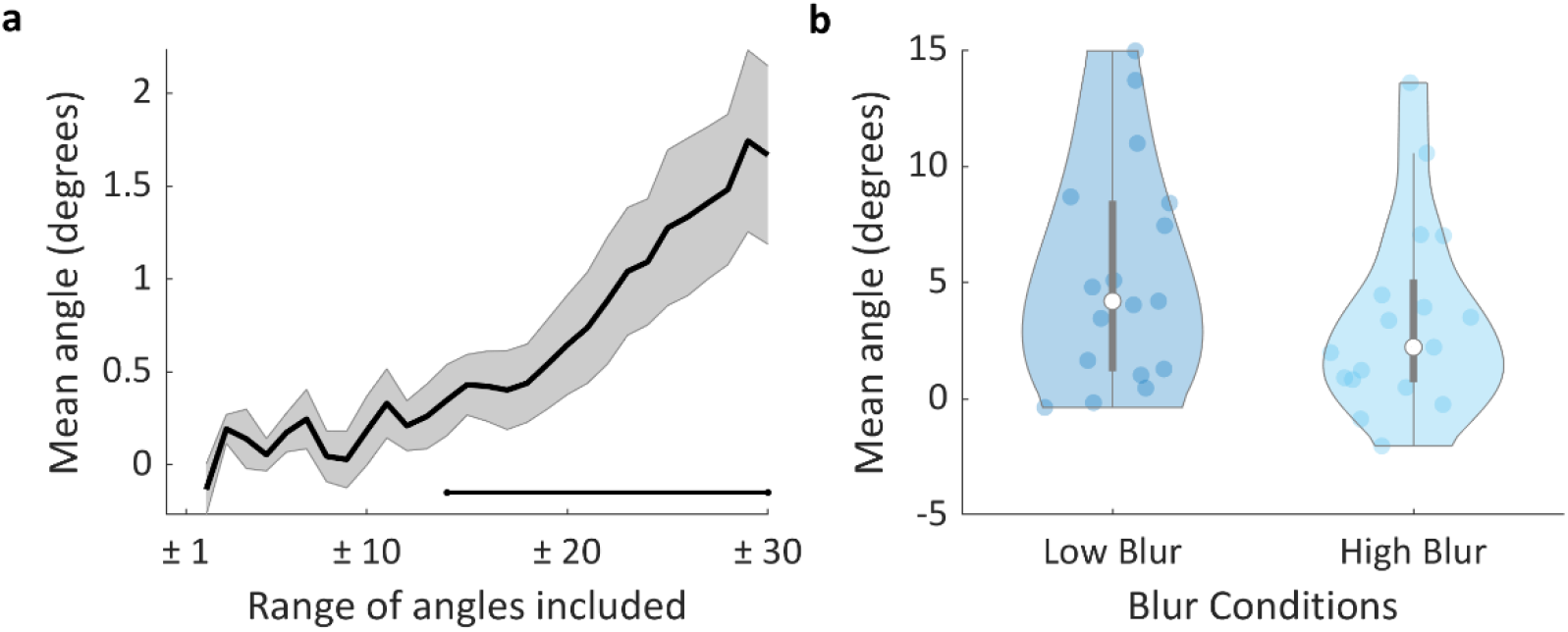
**a.** Mean initial trajectory angle for all blur trajectories (pooled LB and HB data), along incremental ranges of angles symmetric around zero. The solid black line corresponds to the inter-individual mean (the grey area represents s.e.m.). The black horizontal line represents significance (right tailt-test, p < 0.05) against the hypothesis that the mean angle is not larger than zero. **b.** Initial angle of trial trajectories for LB and HB conditions. The coloured dots represent each participant’s mean value for the corresponding condition. The white dots represent the median for each condition and the grey lines illustrate the inter-quartile range.

**Figure 6.**
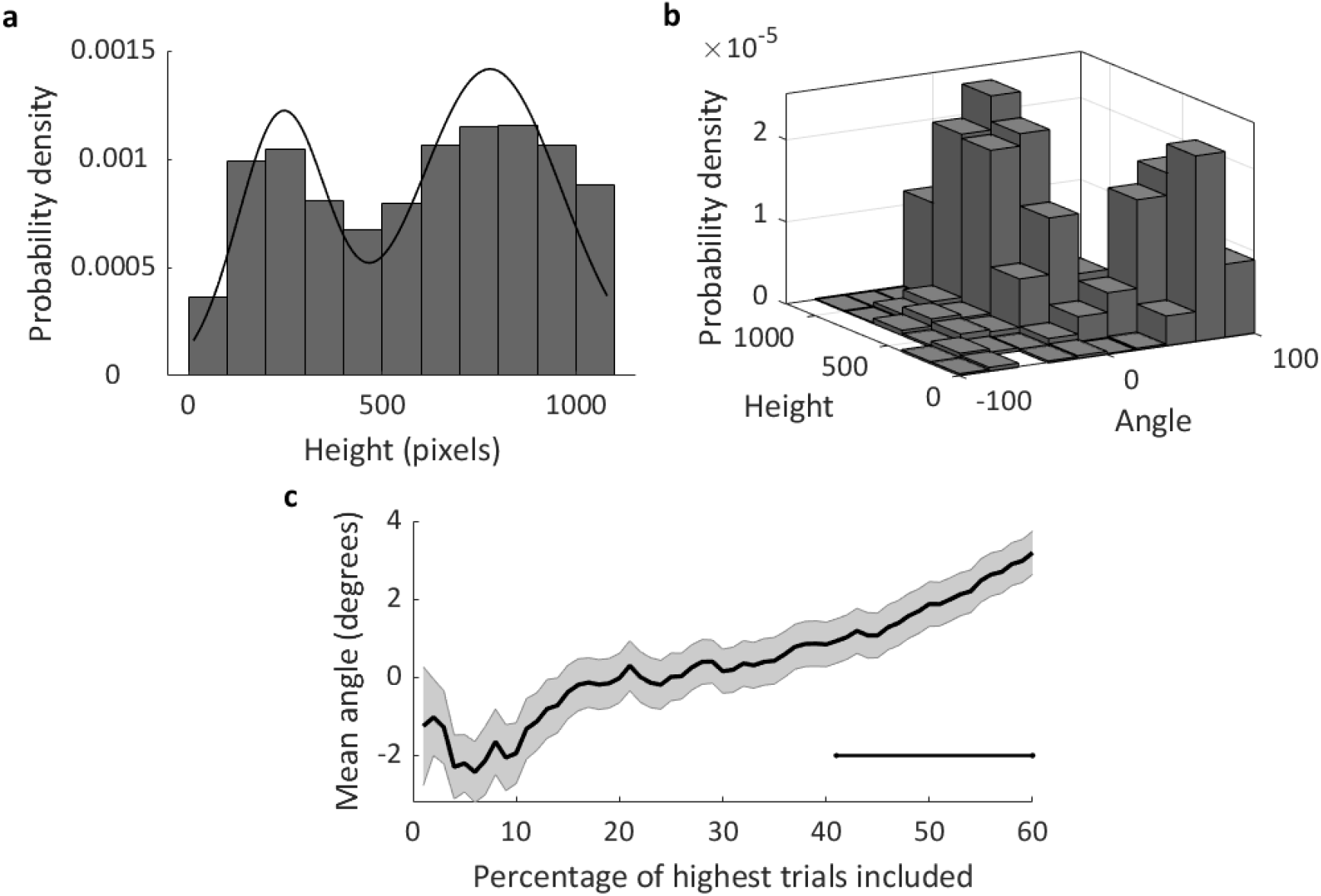
**a.** Probability density of the heights of the trajectories across participants. The solid blackline corresponds to the Gaussian mixture model with 2 components fit to the distribution (better fit in the model with 2 components, AIC = 26439 lower than the model with 1 component, AIC = 26874). **b.** Probability density of the heights and angles of the trajectories across participants. **c.** Mean angle for all blur trajectories (pooled LB and HB data), along incremental ranges of heights. The solid black line corresponds to the inter-individual mean (the grey area represents s.e.m.). The black horizontal line represents significance (right tail t-test, p < 0.05) against the hypothesis that the mean angle is not larger than zero. Angle 0° corresponds to pure vertical upwards movement, i.e., with no horizontal component.

Although we did find those significant deviations in the initial angle of sampling trajectories, we did observe only marginal evidence that the angle deviation was larger in LB (mean = 5.28°, sd = 4.75) than in HB (mean = 3.42°, sd = 4.13) conditions (Fig. 5b; right tail paired-samples t-test, t(16) = 1.66, p = 0.058, Cohen’s d = .4).

### 3.3. Converging evidence from angle and height information

Initially we had decided to classify sampling and non-sampling trials based on initial angle of the trajectories. However, if our hypothesis is correct, a similar classification should apply to the heights of the trajectories. This is because sampling trajectories are expected to reach higher than non-sampling trajectories, as the latter correspond to ballistic movements to the target without much ongoing deliberations and thus are expected to reach vertically much lower. What is more, if trajectories are truly separable into sampling and non-sampling, then it should be the case that in their heights should also be distributed in a bimodal way, and height and angle should be correlated. Consistent with this prediction, we found that heights were distributed in a bimodal way (Figure 6a) across conditions and participants (Figure 6a; Hartigan’s Dip Test, p < 0.05; see Figure S3 for each blur condition). These results in turn suggest that it should be possible to classify trajectories as sampling and non-sampling based on the bimodality in heights, and that this classification should be largely consistent with the one derived above from the angle analyses. In line with this, classification based on height and classification based on angle were highly correlated (Pearson’s correlation, r = 0.76) and clustered trials in two clear categories (Figure 6b).

Similar to the angle analysis reported in Section 3.2 (where trial classification was based on angle), we analysed angle again but this time using trial classification based on height. We found that the angles in sampling trials, both the LB and the HB conditions, were significantly larger than zero (right tail one sample t-test, t(16) = 3.7, p < 0.001, Cohen’s d = .9 and t(16) = 2.05, p = 0.029, d = .5, respectively). This outcome supports the conclusions of our main analysis reported above and shows that this finding generalizes regardless of the classification variable used. Finally, and parallel to the incremental angle analyses reported earlier, we also addressed how the mean angle changes as a function of height increments of trajectories. As one would expect if trajectories reflect both choice and information gathering, we found that as trajectories with lower heights are included into the analysis, mean angle increases (Figure 6c). This shows the interaction between trajectory height and initial angle.

### 3.4. Robustness of the results at earlier initial angles

In the main analysis, we have estimated angles at one third of the trajectory, as we wanted to capture the initial moments of the movements. However, the criterion to compute angles at one-third of the trajectory is somehow arbitrary. As a check regarding the trial classification, we decided to re-compute the trajectory angles at an earlier point in trajectory (described in the Results section). The motivation was to provide an additional look at the angle analysis to reveal that it is robust even at earlier moments of the trajectory. This time we looked at angle at the one-fifth of trajectory as opposed to one-third of trajectory point (described in Figure 2a). The distribution of angles in this calculation also brought about a strong bimodality (Hartigan’s Dip Test, p < 0.05), confirming the main findings. Then, we also corroborated that the angles in blur conditions were significantly larger than zero even at this earlier point. In the LB condition the angle deviated significantly above zero (vertical), but in the HB condition they were not significantly larger than 0 (right tail one-sample t-tests, t(16) = 3.37 p = 0.002, Cohen’s d = 0.81 and t(16) =-0.087, p = 0.53, d =-.02, respectively). This means that whereas the decision starts to have an impact earlier on in the trajectories of LB conditions, in the HB condition the effect is weaker as more information is needed. Moreover, investigating angular deviation at incremental ranges of angles, we found that the angles differed significantly from zero from 22° onward (right tail t-tests, p <.05, Figure S4a). Finally, we repeated the comparison between LB (M = 2.9, sd = 3.57) and HB (M =-.08, sd = 4.001) which showed a strong evidence for the effect of information sampling requirements on the decision status (Figure S4b, right tail paired samples t-test, t(17) = 2.51, p = 0.011, Cohen’s d = 0.61). This additional analysis, calculating the angles from one-fifth of trajectories, provides more confidence regarding the difference between LB and HB conditions, in support for ECT. Based on these converging results, we found strong evidence supporting that there was an impact of the decision component in trajectories overall in the sampling trajectories and that, if any, differences between blur conditions leaned in the expected direction.

## 4. Discussion

Many studies in the past have challenged the classical view of decision making and cognition which assumes a temporal and functional separation between decision and action systems (Pylyshyn, 1984). The idea is that natural choice behaviours of humans and other animals involve movement patterns that reflect the ongoing decision process. As a result, movement trajectory analyses are increasingly used to trace the underlying decision dynamics. Our study clearly sides with these findings, showing that it is possible to trace decision dynamics from the ongoing choice action (Tabor, et. al., 1997, Magnuson, 2005, Spivey & Dale, 2006). However, the tasks used in previous studies did not contemplate the case where actions are also needed to sample information. To fill this gap, we took one step forward from this past research and tested whether the decision outcome pervades action when information sampling is necessary. This is a condition that characterises choice in many natural environments, such as getting closer to an object to decide whether it is food or not.

As mentioned in the introduction, parallel processing of decision-making and action control is an important principle. However, the nature of the interaction between the two is still under debate. For instance, Lepora & Pezzulo (2015) have put forward the ‘embodied choice’ framework, that accommodates richer interactions between action and decision through action-dependent information gain, compared to the parallel account. The experimental tasks they had used to illustrate their predictions lacked the active sampling component, which leaves one main prediction of the theory still unresolved. Our findings support the ‘embodied choice’ theory by showing that decision and action interaction can be traced in ecological scenarios incorporating the active sampling constrain. If this were not the case, we would have observed a temporally separated sampling and responding characteristics in the movement trajectory without any angular deviation in early parts of the trajectories.

One central feature of the task used in the present study is that participants have to trade off information (image de-blurring) for energetic efficiency (moving up, hence orthogonal to the choice goal). This is because motor execution involves expenditure of energy, thus incurring effort-related costs. Motor cost and physical effort have been started to be studied in relation to decision making (Burk, et. al., 2014, Marcos, et. al., 2015). For instance, Cos, et. al. (2014) have shown that effort and biomechanics of a task influence the decision dynamics starting at early stages. It is likely that physical effort influences the decision dynamics due to the strong interactions between action and decision. In our experiment, each blur condition had a different cost/information structure. Although, it is not easy to quantify exactly how this effort to information ratio impacted our results (due to the use of real images instead of parametric stimuli), it is still safe to say that associated effort to sample information altered the decision making process and led to different choice trajectories. The analyses showing an inverse relationship between image visibility and trajectory height clearly support this.

The main result to emerge from this study, however, was based on the deviations and curvatures in choice trajectories. Please note that this is superficially similar to many other mouse-tracking studies (Spivey,et. al., 2010, Freeman, 2018, Wojnowicz, et. al., 2009). A common task characteristic our current study shares with this previous work is the urgency of responding to a task (Scherbaum & Kieslich, 2008. Kieslich, et. al., 2019). Via imposing time pressure, participants are encouraged to execute decision and action in the same time window as it is more optimal for a successful response than waiting statically to make a decision and then move to report it. However, the fundamental difference between our experiment though is the functional link between information and movement. In those previous works, the subject planned and performed actions to report the choice response, therefore effectively allowing to study interactions between decision process and response plan only (as shown in Figure 1a). In contrast, the task we developed here involves, and makes it possible to study, both response and sampling plans and their interplay (Figure 1b). Another way to put it is that previous studies so far have considered only tasks equivalent to the ‘no blur’ condition in of our study. Hence, one of the main goals here was to compare the trajectories between different sampling conditions as a function of movement-information ratio. First, the results obtained conclusively support the prediction that the decision process pervades information sampling movements in various ways. Information sampling trajectories deviated to one of the choices (the correct one, on average) very early on. We confirmed this both in low blur and high blur conditions, using only trials classified as sampling trials. A second expectation by hypothesis was that, if the sampling component was stronger in HB than in LB, then one would assume that the decision component will be more pronounced in LB than in HB trajectories, especially at the early stages. This is because the need for information in HB trials is stronger. Angular differences between LB and HB conditions calculated according to the planned analysis (at 1/3th of trajectories) were in the expected direction, but reached only a marginally significant effect. This borderline result may be due to the fact that the two conditions were not sufficiently different in terms of costs of sampling movement. This cost depended directly on the blur function, which was chosen arbitrarily. Indeed, subsequent analyses where angle was calculated at a more initial stage (1/5th of trajectories), or when angular deviation was calculated in incremental steps from movement origin, revealed robust significant differences in the same, expected direction. This variability reflects the importance of the task mechanics to the study of sensorimotor interactions in a decision making setting (Scherbaum & Kieslich, 2018). Variants of active sampling decision making tasks, including variations of the information cost function, should shed more light on the full range of embodied decisions under naturalistic constrains.

The proposed interactions between action and decision we suggest rely on the incorporation of sampling and responding actions in the task structure (Figure 1b). We note that the tasks that include movement-agnostic stimulus, often used in the literature (and summarised in Figure 1a), are a special instance of the more general case modelled in Figure 1b: one in which the arrows to and from “sampling plan” have zero weight. Yet, our experimental setup is not intended to as a general model for all action-decision possibilities that humans and animals are capable of. We rather claim that embodied decisions are the manifestation of the flexibility of the decision process (Wispinski, et. al., 2018). In many natural and ecological situations, like the one modelled here, decisions have to be carried out as ETC predicts –with a strong interaction coupling with action processes. Nevertheless, there are also abstract and higher-level decisions which may comply with serial accounts of decision making, especially in humans. In line with a ‘phylogenetic refinement’ view, fully abstract cognitive operations are evolutionarily more recent, whereas rich cycles of action & decision are prevalent from very basic animals to complex mammals (Cisek, 2019). In the human context, depending on the task, the biomechanical characteristics and previous experience, we may observe response patters ranging from a pure abstract and covert decision making process that precedes any action, to a fully embodied and interactive one such as the one seen here. For instance, a novice driver may find herself thinking step-by-step about all of the driving actions before executing them, however as practice accumulates, she may decide and move at the same time with ease. Therefore, we are aware of the vast complexity about the interaction between decision and motor action (Gallivan, et. al., 2018). Our study provides a step forward in understanding these interactions under the new constrain of action-dependent information sampling. What we have shown is that when the task dynamics imposes this type of ecological constraint, action for sampling and choice action have interactions with the decision process and with each other.

To summarize, here we showed a demonstration of interactions between action to sample information, action to respond and decision process with a novel mouse-tracking task. Our results showed that decision feed into movement trajectory during information sampling movements. This is a support for the embodied theories in decision making in a way that has been lacking in the field, as far as we know.

## 5. Acknowledgements

This project receives support from the grant from Agency for Management of University and Research Grants (AGAUR-2017 SGR 1545), Ministerio de Economia y Competitividad (PSI2016-75558-P AEI/FEDER). This project has been co-funded with 50% by the European Regional Development Fund under the framework of the ERFD/FEDER Operative Programme for Catalunya 2014-2020.

D.O. is supported by a fellowship from “la Caixa” Foundation (ID100010434; fellowship code LCF/BQ/DI18/11660026). This project has received funding from the European Union’s Horizon 2020 research and innovation programme under the Marie Skłodowska-Curie grant agreement No. 713673.

RM-B is supported by BFU2017-85936-P from MINECO (Spain), the Howard Hughes Medical Institute (HHMI; ref 55008742), an ICREA Academia award, and the Bial Foundation (grant number 117/18). S.S-F. is funded by Ministerio de Ciencia e Innovación (Ref: PID2019-108531GB-I00 AEI/FEDER).

## 6. Supplementary Figures

**Figure S1.**
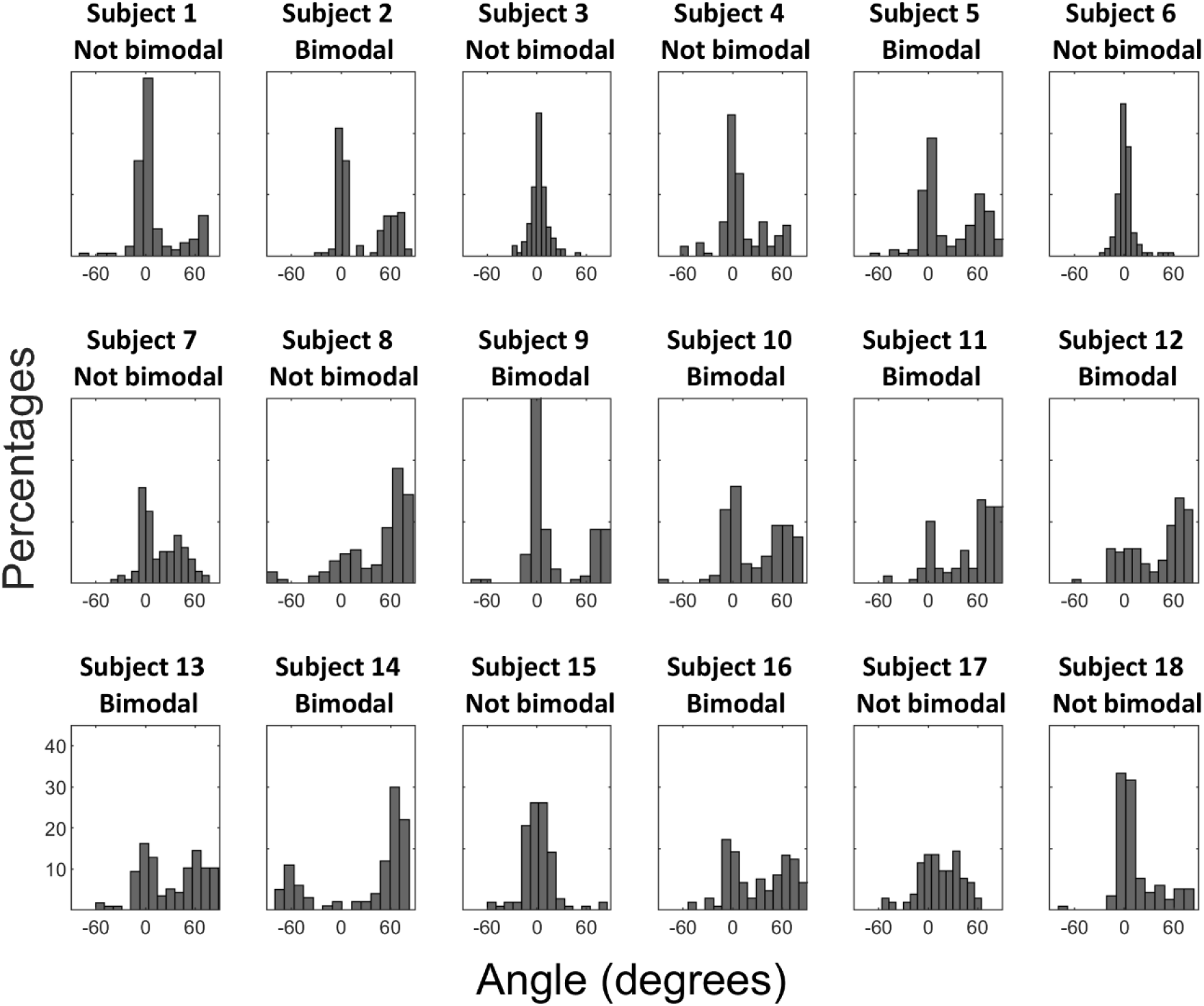
Distribution of angles for each individual subject.

**Figure S2.**
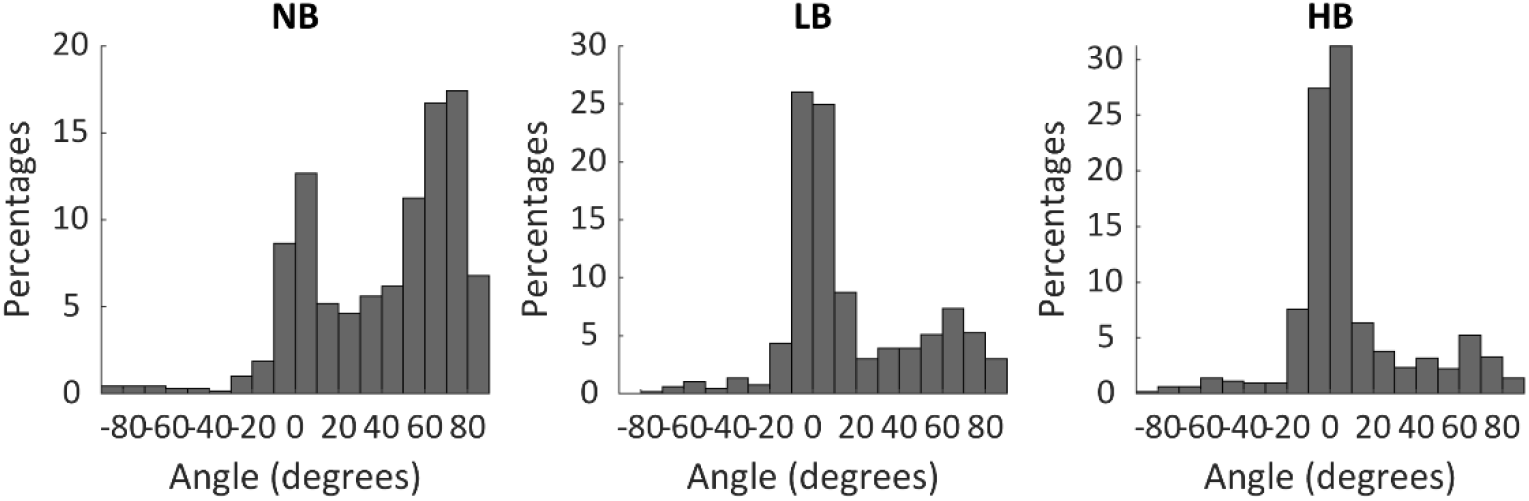
Distribution of trajectory angles for each blur condition.

**Figure S3.**
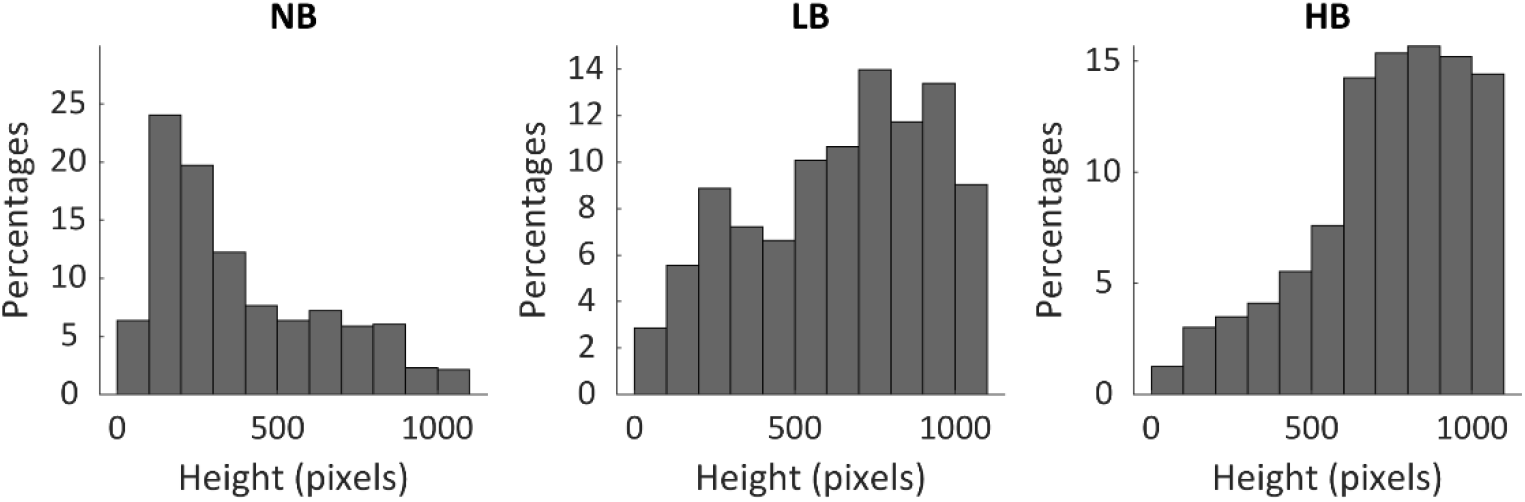
Distribution of trajectory heights across participants for each blur condition.

**Figure S4.**
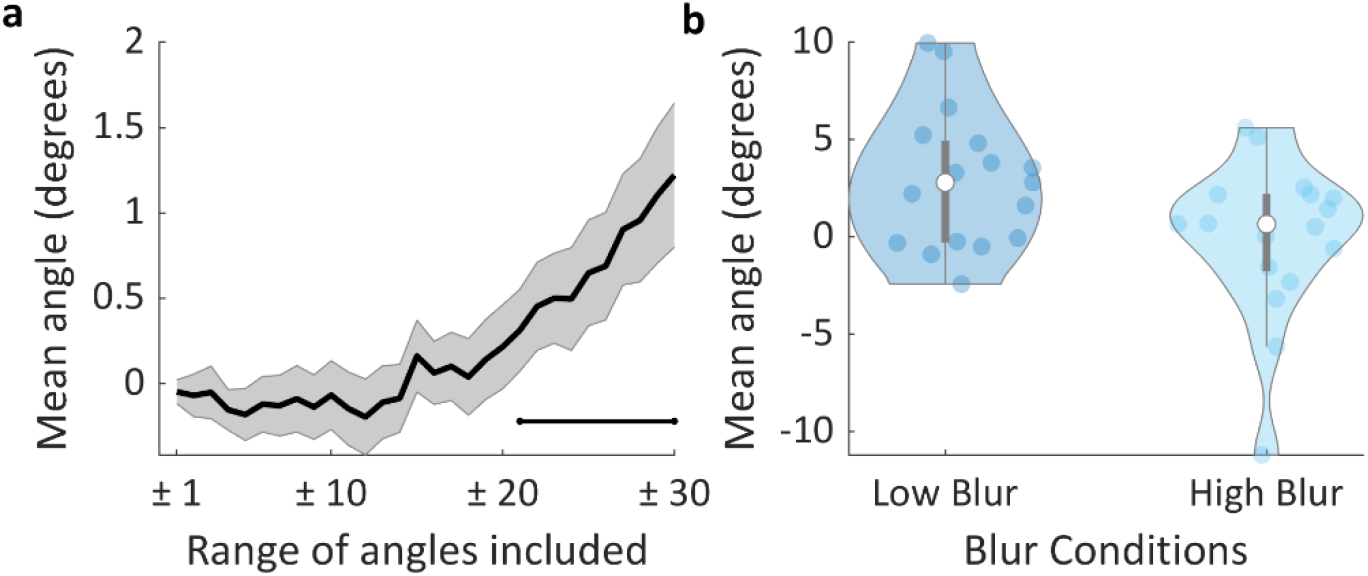
Analysis on angles which was calculated based on one fifth of trajectory length **a.** Mean angle for LB & HB trajectories, for different ranges of angles, symmetric around zero, included in the analysis. Full line corresponds to the mean; grey area represents s.e.m across subjects. The black horizontal line represents significance (Right tail one-sample t-test, p < 0.05) against the hypothesis that the mean angle is not larger than zero. **b.** Mean initial angle of trial trajectories for LB and HB conditions. The grey lines represent each participant’s mean value for the corresponding condition. The dark line is the sample mean of all data, error bars representing s.e.m.

## 7. Supplementary Table

**Table S1.** Bayesian counterparts of the t-tests that have been reported in the Results section. The analyses are ranked in the order of appearing in text.

|  | Page | Analysis Variable | H1 | Bayes Factor | Error | Median Effect Size | 95% CI | Frequentist p-values |
| --- | --- | --- | --- | --- | --- | --- | --- | --- |
| 1 | 7 | Height (all trials) | HB > LB | 1067.6 | < 0.001 | 1.17 | [0.54, 1.79] | <0.001 |
| 2 | 8 | Angle (classification based on angle) | LB > 0 | 209.53 | <0.001 | 1.01 | [0.41, 1.63] | <0.001 |
| 3 | 8 | Angle (classification based on angle) | HB > 0 | 25.27 | <0.001 | 0.73 | [0.21, 1.29] | =0.002 |
| 4 | 9 | Angle (classification based on angle) | LB > HB | 1.45 | 0.004 | 0.37 | [0.04, 1.12] | =0.058 |
| 5 | 11 | Angle (classification based on height) | LB > 0 | 44.02 | <0.001 | 0.8 | [0.26, 1.38] | <0.001 |
| 6 | 11 | Angle (classification based on height) | HB > 0 | 2.56 | 0.002 | 0.44 | [0.06, 0.93] | =0.029 |
| 7 | 11 | Angle (calculated at one-fifth) | LB > 0 | 22.2 | < 0.001 | 0.72 | [0.2, 1.27] | =.002 |
| 8 | 11 | Angle (calculated at one-fifth) | HB > 0 | 0.23 | 0.004 | 0.143 | [0.01, 0.49] | =.53 |
| 9 | 12 | Angle (calculated at one-fifth) | LB > HB | 5.36 | <0.001 | 0.54 | [0.1, 1.05] | =0.011 |

